# *Theileria annulata* histone deacetylase 1 (TaHDAC1) initiates schizont to merozoite stage conversion

**DOI:** 10.1101/2022.06.20.496823

**Authors:** Shahin Tajeri, Laurence Momeux, Benjamin Saintpierre, Sara Mfarrej, Alexander Chapple, Tobias Mourier, Brian Shiels, Frédéric Ariey, Arnab Pain, Gordon Langsley

## Abstract

A fungal metabolite, FR235222, specifically inhibits a histone deacetylase of the apicomplexan parasite *Toxoplasma gondii* and TgHDAC3 has emerged as a key factor regulating developmental stage transition in this species. Here, we exploited FR235222 to ask if changes in histone acetylation regulate developmental stage transition of *Theileria annulata*, another apicomplexan species. We found that FR235222 treatment of *T. annulata*-infected transformed leukocytes induced a proliferation arrest. The blockade in proliferation was due to drug-induced conversion of intracellular schizonts to merozoites that lack the ability to maintain host leukocyte cell division. Induction of merogony by FR235222 leads to an increase in expression of merozoite-marker (rhoptry) proteins. RNA-seq of FR235222-treated *T. annulata*-infected B cells identified deregulated expression of 468 parasite genes including a number encoding parasite ApiAP2 transcription factors. Thus, similar to *T. gondii*, FR235222 inhibits *T. annulata* HDAC (TaHDAC1) activity and places parasite histone acetylation as a major regulatory event of the transition from schizonts to merozoites.

## Introduction

Bovine tropical theileriosis due to leukocyte infection by *Theileria annulata* is a disease of considerable importance across several continents. Acute infection in susceptible hosts results in mortality in less than a month and sub-acute infection impedes weight gain and milk production. A huge population of cattle are at risk of infection in disease endemic areas^1^. The obligatory blood feeding behaviour of ticks of the genus *Hyalomma* is mainly responsible for parasite transmission to cattle. Tick-derived *T. annulata* sporozoites infect bovine leukocytes and develop into a poly-nucleated cell termed the (macro) schizont^2^. The schizont stage is able to transform its host leukocyte into an immortalized cancer-like cell that disseminates throughout the animal^3^. At some point, schizonts increase their nuclei number and develop via a process called merogony into single cell extracellular merozoites that rapidly invade red blood cells. Inside red blood cells, merozoites become small, pear-shaped piroplasms, hence tropical theileriosis is also known as piroplasmosis. Piroplasms circulate in red blood cells and are acquired by the tick vector during blood feeding. A phase of sexual reproduction then occurs, with the tick phase of the life cycle culminating in the production of haploid sporozoites in the salivary gland. Sporozoites are then transmitted to cattle during the blood meal of the nymph or adult tick^2^. A hydroxynapthoquinone derivative, buparvaquone is widely used to treat clinical disease, but parasite resistance to the drug is known to be spreading rapidly^4, 5^. Long-term passage schizont-infected transformed leukocytes are used as live attenuated vaccines against tropical theileriosis, and together with buparvaquone and intensive tick control measures constitute the major strategies to control disease^6^.

In the context of the *T. annulata*-infected bovine leukocytes, virulence has two major components. First and foremost, is the ability of the intracellular schizont to transform host myeloid cells and B lymphocytes^7^. Through secretion of a repertoire of proteins on to the parasite surface or translocated into the host cell compartment, the schizont fine tunes host cell signalling pathways to create an infected leukocyte with cancer-like properties^8^. *T. annulata*-transformed leukocytes have high dissemination potential and secrete a range of proteases and inflammatory cytokines that influence their pathogenicity^9^. Among these proteases is matrix metalloproteinase 9 (MMP9)^10^ known to mediate malignancy of several human and animal tumours^11^. Activating protein 1 (AP-1) is a heterodimeric transcription factor induced by *Theileria* infection and it drives transcription of *mmp9*^12^. Pharmacological inhibition of MMP9 blocks dissemination of *Theileria-*transformed B cells and macrophages in immunodeficient mice^13^. Parasite dependent activation of a host enzyme c-Jun N-terminal kinase (JNK) contributes to AP-1 activity through direct phosphorylation of c-Jun (an AP-1 family member)^14^. Infection also upregulates a host methyltransferase (SMYD3) that methylates bovine histone 3 at lysine 4 to maintain active transcription of *mmp9*^15^. The other important component of *T. annulata-*infected leukocyte virulence is the ability of the intracellular schizont to produce infectious merozoites that rapidly invade red blood cells, becoming piroplasm infected RBC that cause significant anaemia^16^. Anaemia clinically distinguishes tropical theileriosis from East Coast Fever (ECF) caused by *T. parva*^17^. In long-term passaged *T. annulata*-transformed cell lines, widely used as live vaccines, both schizont and merozoite-associated pathologies are severely dampened^18, 19, 20^. Thus, schizont-induced transformation of host leukocytes and merozoite-infection of red blood cells are the two major events of *T. annulata* infection that cause clinical disease.

The ability of two distinct *T. annulata* developmental stages with highly specialized adaptations to survive inside either a nucleated (leukocyte) or non-nucleated (erythrocyte) mammalian cell is likely to be determined by their stage-specific gene expression programs. While knowledge on mechanisms underlying stage-specific gene regulation in closely related apicomplexan parasites *Plasmodium* and *Toxoplasma* is increasing at a high pace, little is known about how *Theileria* parasites control expression of their genes. However, similar to other Apicomplexa, the *Theileria* genome encodes a number of Apicomplexa Apetala-2 type (ApiAP2) transcription factors, but except for one report^21^, these transcription factors have not been studied in great detail. The limited number of ApiAP2 family of genes in apicomplexan genomes (20 genes in *T. annulata*, 27 genes in *Plasmodium* and 67 members in *T. gondii*) and the extensive phenotypic plasticity of their complex life cycles suggests extensive epigenetic control of gene expression operates. In support of this, a plethora of epigenetic modifiers and DNA-interacting proteins have been discovered in apicomplexan genomes. Epigenetic gene regulation involves modulation of chromatin structure through posttranslational modification (PTM) of histone proteins without changes in nucleic acid sequences. Epigenetic modifier enzymes of note are histone acetyltransferases (HATs), histone deacetylases (HDACs), methyltransferases and demethylases^22^. HAT-catalyses transfer of an acetyl group to histone tails resulting in relaxation of heterochromatin providing transcription factor accessibility sites to promote and maintain gene expression. HDACs counteract HAT function and are capable of supressing gene expression^23^. Both HATs and HDACs can also modify non-histone substrates leading to changes in protein localization, stability^24, 25^.

The genetic tractability of *Plasmodium* and *Toxoplasma* has led to detailed characterization of some of these enzymes and their importance in stage-specific gene expression. For instance, TgGCN5 (an acetyltransferase) has been found to acetylate histone 3 at lysine 18 (H3K18ac) and TgCARM1 (a methyltransferase) methylates arginine 17 of H3 (H3R17me)^26^. Interestingly, pre-infection exposure of tachyzoites to a specific inhibitor of TgCARM1 promoted formation of intracellular cysts (containing bradyzoites) post invasion^26^. In *T. annulata* a methyltransferase (TaSETup1) deposits methyl groups on H3K18 to repress gene expression, and drug-mediated inhibition of demethylation dampened development of merozoites in heat stressed schizonts^27^. Thus, similar to other Eukaryotes, apicomplexan epigenetic modification enzymes seem to have seminal roles in their infection biology, and chemical or genetic ablation of their activities causes parasite life cycle arrest, or promotion/blockade of stage differentiation. General histone deacetylase inhibitors (HDACi) apicidin and TSA are able to block constant re-initiation of intraerythrocytic schizogony of *Plasmodium* in red blood cells^28, 29^ with TSA possessing strong gametocytocidal activity^30^. In addition, they can eliminate *Toxoplasma* tachyzoites and induce tachyzoite to bradyzoite conversion^31^. Interestingly, a cyclopeptide fungal metabolite with structural similarities to apicidin, FR235222, can penetrate the cell wall of *Toxoplasma gondii* cysts and affect dormant bradyzoites^32^. FR235222 was initially isolated from the fermentation broth of *Acremonium* species^33^ and showed activity against *T. gondii* and *P. falciparum* parasites, and its specific target molecule was found to be *T. gondii* HDAC3 (TgHDAC3)^31^.

Given the presence of a highly conserved orthologue of TgHDAC3 in *Theileria* parasites (herein named TaHDAC1, like PfHDAC1), availability of an apicomplexan-specific inhibitor and the pivotal role of TgHDAC3-associated proteins in *Toxoplasma* life cycle progression, we decided to examine the potential role of TaHDAC1 in regulating *T. annulata* developmental progression. We report that drug mediated inhibition of a TaHDAC1 at the schizont stage initiates a switch in development to the merozoite stage. In this manner, the function of TaHDAC1 as a regulator of differentiation in *Theileria* and *Toxoplasma* appears conserved. Identification of *T. annulata* genes with modulated expression induced by FR235222, notably included 5 upregulated TaApiAP2 transcription factors. Our results provide a better understanding of how initiation of merogony is regulated in this fascinating model of animal parasitism, and supports the notion that a basic mechanism of stage differentiation operates across the Apicomplexa.

## Results

### FR235222 blocks proliferation of *Theileria annulata*-transformed B lymphocytes and macrophages

Exploiting the conserved nature of histone deacetylase domains, we queried the *T. annulata* reference genome (Ankara C9)^34^ and identified four genes predicted to harbor histone deacetylase domains (**Fig. 1a**). One of these putative histone deacetylases (TA12690) showed a high degree of similarity with *Toxoplasma gondii* HDAC3 (TgHDAC3) and with HDACs of other apicomplexan parasite species (**Supplementary Fig. 1**). Since all of these apicomplexan orthologues (except TgHDAC3) have been annotated as histone deacetylase 1, we named TA12690 as *T. annulata* histone deacetylase 1 (TaHDAC1). In addition, the availability of a specific inhibitor of TgHDAC3 prompted us to examine the role of TaHDAC1 in *T. annulata*-infected leukocytes. Treatment of infected cells with FR235222 significantly dampened proliferation of *T. annulata-*infected bovine B cells (TBL20) and macrophages, while not affecting proliferation of uninfected BL20 B cells (**Fig. 1b, c** and **Supplementary Fig. 2**). The proliferation arrest observed for FR235222-treated leukocytes was not due to host cell death, as more than 90% of infected leukocytes were viable following 48-72 h of treatment (**Supplementary Fig. 3**). Eosin/Azur-stained smears revealed an evident increase in parasite nuclei number per infected cell following FR235222 treatment. (**Fig. 1d**). The increase in nuclei number was confirmed by immunofluorescence, where intracellular parasites were detectable by staining with an antibody specific marker (TA08425=p104) and DAPI (**Fig. 1e**). Further manual counting of parasite nuclei revealed the increase to be significant (**Fig. 1f**). Importantly, FR235222 treatment induced hyperacetylation of parasite histone H4, as previously reported in *T. gondii*^*31*^ (**Supplementary Fig. 4**), consistent with the drug blocking TaHDAC1 activity. Therefore, FR235222-mediated inhibition of histone deacetylation by TaHDAC1 correlates with a drop in *T. annulata*-induced leukocyte proliferation. Moreover, as it is known that an increase in parasite nuclear number precedes commitment to merozoite production we postulated that blocking TaHDAC1 activity promotes initiation of schizont to merozoite differentiation.

**Fig. 1:**
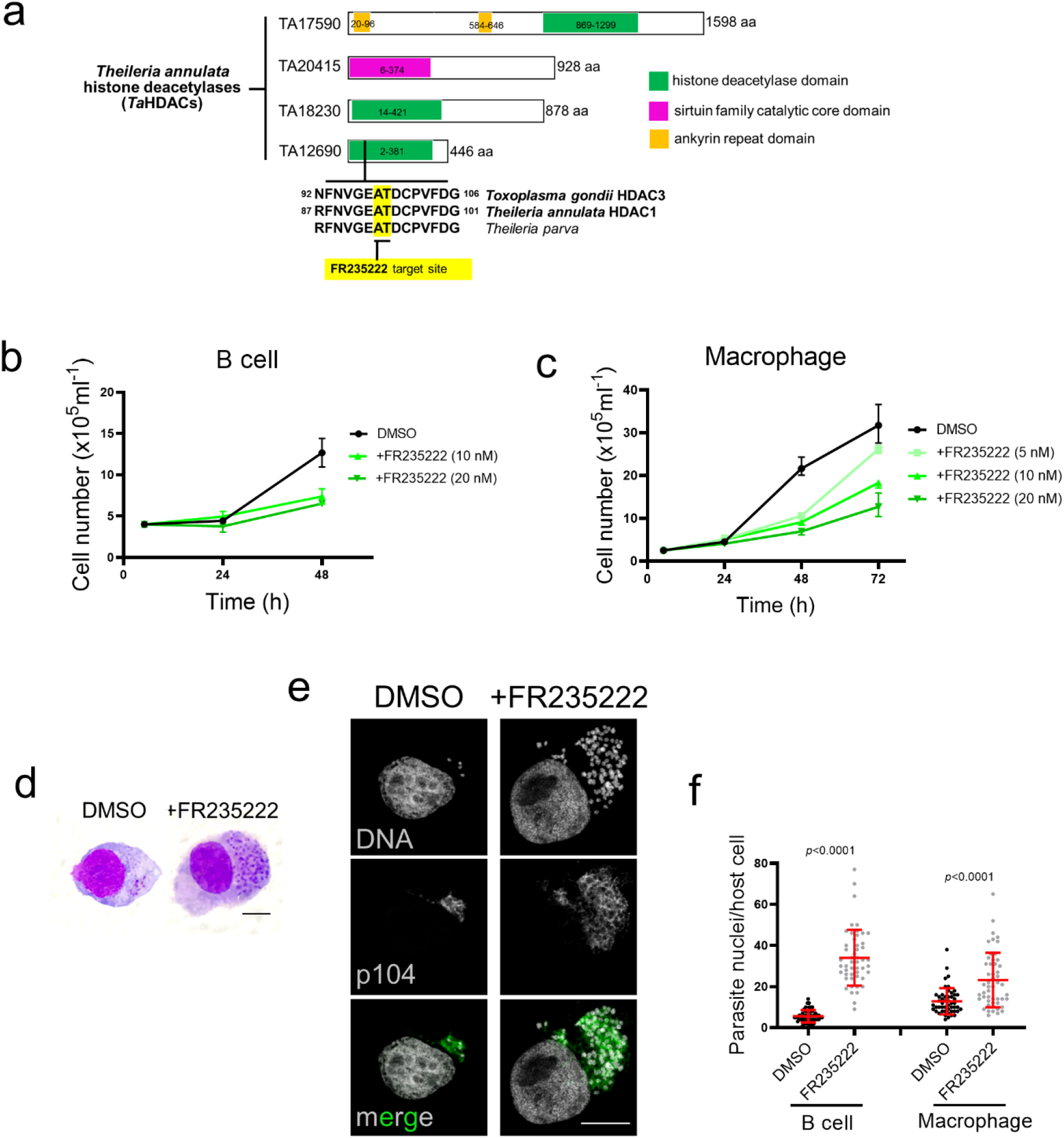
FR235222 inhibition of *Theileria annulata*-transformed leukocyte proliferation. **a** Schematic representation of the four histone deacetylase enzymes encoded by the *T. annulata* genome and their domain(s) organization. The genes are identified by their Gene IDs at PiroplasmaDB. The conserved FR235222 target site originally identified in *Toxoplasma gondii* HDAC3 and located within the deacetylase domain is highlighted in yellow. Amino acid = aa. **b** Proliferation profiles of *T. annulata*-transformed B lymphocytes (TBL20); **c** Ode macrophages. Both infected B cells and macrophages exposed to increasing doses of FR235222. Proliferation curves represent a typical example of several reproducible experiments. **d** Light microscopy images of Eosin/Azur stained preparations from control and FR235222-treated parasitized B cells, 100X objective lens with oil, bar = 10 μm. **e** Immunofluorescence images of TBL20 B cells treated with 10 nM FR235222 compared to DMSO-only exposed TBL20. Host and parasite DNA labelled with DAPI and parasites detected by monoclonal antibody (1C12) specific for the schizont surface p104 protein. Scale bar = 10 μm. **f** Quantification of schizont nuclei counted in 50 individual infected leukocytes (TBL20 and Ode macrophages) stained with DAPI and observed under an immunofluorescence microscope. Two-tailed Student’s t-test was used to estimate significance.

### FR235222-induced blockade of *T. annulata*-induced leukocyte proliferation is due to parasite developmental stage differentiation

We next asked if the dampening in proliferation following 48h of FR235222 treatment is due to the transforming schizont stage differentiating to non-transforming merozoites, a process termed merogony. Induction of merogony is considered stochastic in *T. annulata*^35^, and can be induced by culturing infected leukocytes for 4-6 days at 41°C^36^. Merogony involves an increase in parasite DNA due to multiplication of nuclei, budding and liberation of single cell merozoites from the schizont syncytium, host leukocyte proliferation arrest and eventual rupture^37^. To assess merogony potential, we prepared mRNA from FR235222-arrested TBL20 infected B cells and verified by qRT-PCR expression of a selected panel of 30 *T. annulata* genes. This panel included: 13 genes coding for proteins orthologous to *T. gondii* proteins, whose chromatin of the corresponding genes was hyperacetylated following FR235222 treatment. The hyperacetylation of their chromatin led to the genes being expressed by non-proliferating bradyzoites and/or sporozoite stages^31^. In addition, as a negative control 17 genes chosen at random were included (**Supplementary Table 1**). Exploiting the limited expressed sequence tag (EST) data available at PiroplasmaDB, we regrouped genes based on EST evidence for merozoite and piroplasm expression. This revealed upregulation of several merozoite-specific genes such as Tamr1 (TA16685) and TaAMA1 (TA02980), plus the genes coding for proteins with homology to the *Toxoplasma* FR235222-target ‘up’ group (**Fig. 2a** and **Supplementary Table 1**).

**Fig. 2:**
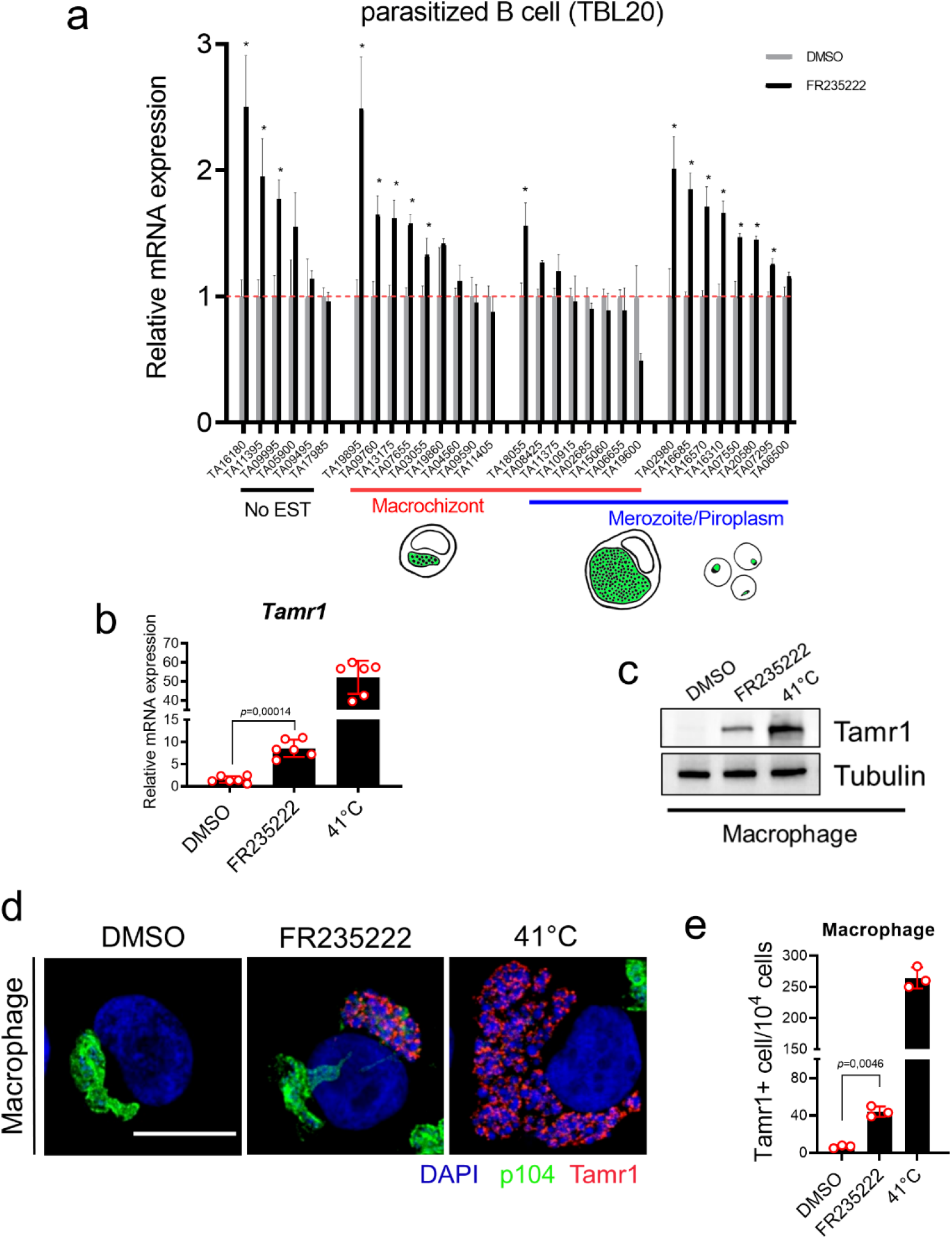
Induction of merogony by FR235222 in *T. annulata*-transformed leukocytes. **a** qRT-PCR amplicons from a panel of 30 parasite genes in TBL20 B cells treated or not with 10 nM FR235222 for 48 h. Genes were grouped based on the presence of corresponding EST at PiroplasmaDB: no EST (black underlined), schizont (red underlined) and merozoite/piroplasm (blue underlined). Genes displaying significant (Student’s t test, *p* value <0.05) increase in transcript levels upon FR235222 exposure are labelled with an asterisk. Gene mean mRNA expression levels ± standard deviations are displayed. *T. annulata* actin gene (TA15750) was used as housekeeping gene control. Tamr1 mRNA expression **b** and protein levels **c** in virulent (passage 52) Ode macrophages cultured at 37°C (DMSO only), incubated with FR235222 for 7 days, and cultured at 41°C (as a positive control for merozoite production). Tubulin was used as a western blot loading control. **d** Confocal microscopic images of Ode macrophages cultured under different conditions. Host and parasite DNA were stained with DAPI. Schizonts were labelled by monoclonal antibody 1C12 against p104 (in green). Merozoite rhoptry antigen (Tamr1) shown in red. Note the disappearance of schizont marker p104 in cells undergoing merogony with FR235222-treatment and Ode macrophages cultured at 41°C. Photos taken with a X63 objective and 2X zoom, scale bar = 10 μm. **e** Manual quantification of Tamr1-positive cells grown under the different experimental conditions. Results representative of three independent experiments.

Since full merogony takes 7-8 days to complete we cultured both virulent Ode macrophages for 7 days at 41°C and compared them to Ode macrophages treated for 7 days with FR235222 at 37°C. Protein and total RNA extracts were prepared and expression of a specific marker for commitment to merozoite production (*T. annulata* merozoite rhoptry protein 1, Tamr1=TA16685) examined. Both culturing at 41°C, and exposure to FR235222 at 37°C induced expression of Tamr1 in virulent parasites (**Fig. 2b-e**). Thus, we conclude that inhibition of TaHDAC1 activity by FR235222 results in an arrest of infected leukocyte proliferation due to schizonts initiating differentiation towards merogony. As treatment with a general HDAC inhibitor (apicidin) has been shown to perturb expression of several *P. falciparum* ApiAP2 genes^29^, we confirmed that apicidin enhances merogony induced by elevated temperature (41°C), but not at 37°C, in the D7 *T. annulata* infected cell line (**Supplementary Table 2 and Supplementary Fig. 5**). Moreover, it was apparent that treatment with apicidin had no effect on the down regulation of the macroschizont marker, p104 that occurs during merogony (**Supplementary Fig. 5**).

### FR235222-induced inhibition of TaHDAC1 impacts significantly on the transcription of 468 *T. annulata* genes

Since FR235222-induced a merogony-related dampening in proliferation of infected leukocytes, both infected TBL20 and non-infected BL20 cells were treated for just 48 h with 10 nM FR235222, a low dose that dampened proliferation of parasitized, but not uninfected B cells (**Fig. 1b** and **Supplementary Fig. 2**). RNA-seq analyses revealed 468 differentially expressed parasite genes (DEGs) that displayed significant (adjusted *p*-value <0.05, more than two-fold up- or down) expression compared to control DMSO-only treated TBL20 B cells. As expected, upon inhibition of TaHDAC1-mediated histone deacetylation a large proportion of the DEG genes (441 genes, i.e. 94.23 %) displayed upregulated mRNA levels. Surprisingly, increased histone acetylation led to a reduced expression of 27 (5.76 %) genes (**Fig. 3a**). Tables containing full lists of DEGs are provided (**Supplementary File 1**). We next assigned available (PiroplasmaDB) expressed sequence tags (ESTs) to all FR235222-induced genes with fold-increase in expression equal to or greater than Tamr1. Merozoite ESTs could be assigned to 126 genes with an expression equal to or greater than Tamr1 (**Table 1**). A two-sided Fisher’s exact test gave a *p*-value of 0.002765 indicating that FR235222-treatment had induced a significant enrichment for genes expressed in merozoites. Genes displaying FR235222-induced expression are equally well distributed over the four *T. annulata* chromosomes and not confined to specific genomic regions (**Supplementary Fig. 6**). As they included *TA16660* that codes for a protein with significant similarity to *P. falciparum* rhoptry neck protein 5 (PfRON5, PF3D7_081770) we confirmed by qRT-PCR its upregulation in FR235222-treated TBL20 B cells (**Supplementary Fig. 7a**). We could identify other FR235222-upregulated genes potentially involved in merozoite invasion of bovine red blood cells. These include the well-known apicomplexan microneme protein apical merozoite antigen (AMA-1, TA02980), several other rhoptry associated proteins and a thrombospondin type-1 repeat containing protein (TRAP, TA14215). All these proteins are predicted to have signal peptides and/or transmembrane domains indicating possible roles at the host-parasite interface (**Table 2**).

**Fig. 3:**
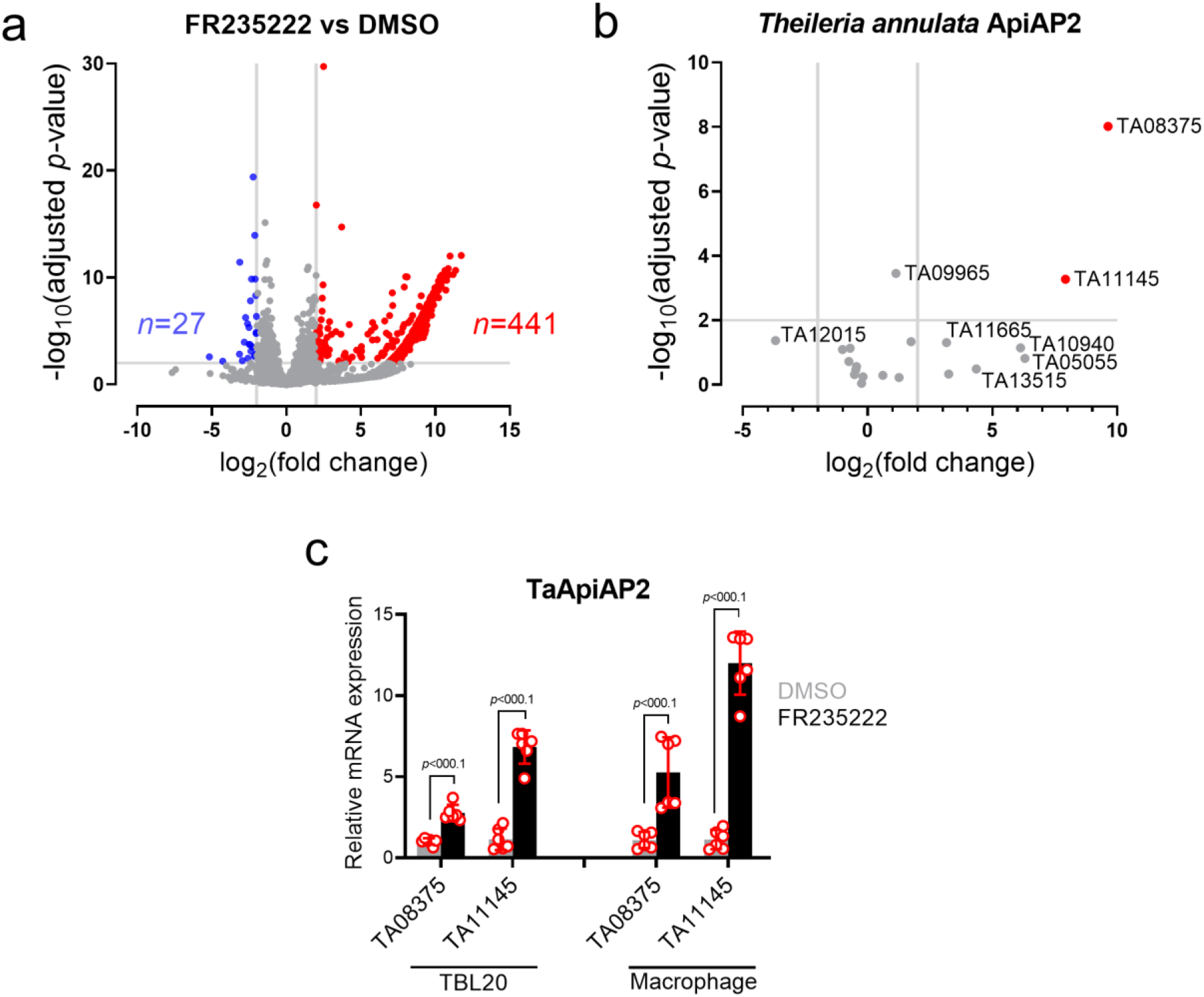
Transcriptional landscape of *T. annulata* following FR235222 inhibition of TaHDAC1. **a** Volcano plot to visualize RNA-seq data of *T. annulata* schizonts (TBL20) treated with FR235222 in comparison to DMSO-only treated TBL20. Each dot represents a parasite gene. Differentially expressed genes (DEGs) that are either significantly higher (red) or lower (blue) in FR235222 treated parasites. Unaffected genes are in grey. Vertical and horizontal grey lines delineate >2 log_2_ fold change and >2 –log_10_ adjusted *p* value (padj), respectively. Volcano plot was generated using GraphPad Prism version 8.4.0. **b** Volcano plot of 20 *T. annulata* ApiAP2 genes. **c** Confirmation of FR235222 induced DEG AP2 genes in TBL20 B cells and virulent Ode macrophages (p52) by qRT-PCR. qRT-PCR data were normalized to TaHSP70 (TA11610). Two-tailed student’s t-test used to estimate significance. Mean fold change in mRNA expression level ± standard deviations shown. qRT-PCR results are representative of two independent experiments.

**Table 1.** ESTs for *T. annulata* genes with expression equal to or greater than TA16685 (Tamr1).

|  | FR235222-induced | Non-FR235222 induced | Total |
| --- | --- | --- | --- |
| **With merozoite EST** | 126 | 560 | 686<br>(*T. annulata* genes with merozoite ESTs) |
| **Without merozoite EST** | 428 | 2682 | 3110<br>(*T. annulata* genes without merozoite ESTs) |
| Total | 554 | 3242 | 3796<br>(Total *T. annulata* genes) |

**Table 2.** Some significant FR235222-induced *Theileria annulata* genes potentially involved in invasion of host cells.

| Gene ID | Product | Description | Fold change | Signal peptide | Transmembrane domain | GPI anchor |
| --- | --- | --- | --- | --- | --- | --- |
| TA02980 | apical merozoite antigen, putative | AMA-1 | +10,14 | - | 1 | - |
| TA21020 | hypothetical protein, conserved | RAP | +9,86 | - | - | - |
| TA20965 | rhomboid family integral membrane protein, putative | Rhomboid protein (ROM) | +8,88 | - | 5 | - |
| TA13245 | hypothetical protein | roptry neck protein 4 (RON4) | +8 | √ | 0 | - |
| TA16660 | hypothetical protein | RON | +7,65 | - | 1 | - |
| TA05705 | roptry-associated protein, putative | RAP-1 | +7,59 | √ | 0 | - |
| TA14215 | hypothetical protein | thrombospondin-related anonymous protein (TRAP) | +7,55 | √ | 1 | - |
| TA03755 | sporozoite surface antigen, putative | SPAG-1 | +7,23 | √ | 1 | √ |

None of the above listed proteins except SPAG-1 has been characterized to date, Thus, their assumed involvement in invasion comes from observations made in other apicomplexan parasites, Presence of a putative signal peptide, or transmembrane domain was estimated by signal-IP v5 (https://services.healthtech.dtu.dk/service.php?SignalP-5,0) and TMHMM (https://services.healthtech.dtu.dk/service.php?TMHMM-2,0) servers, respectively, Presence glycosylphosphatidylinositol (GPI) anchor signal was verified by GPI-SOM (http://gpi.unibe.ch/).

Next, we focused on parasite ApiAP2 transcription factors due to their potential involvement in regulation of *T. annulata* merozoite production^21^. Expression of six different TaApiAP2s was altered following FR235222-induced inhibition of TaHDAC1, but only two (*TA08375* and *TA11145*) were DEG at 48 h (**Table 3 and Fig. 3b**). FR235222 mediated upregulation of *TA08375* and *TA11145* was confirmed in transformed B lymphocytes and macrophages by qRT-PCR (**Fig. 3c**). Four other ApiAP2s (*TA13515, TA16485, TA05055* and *TA10940*) showed a non-significant (>3 log fold-change) increase in expression by RNA-seq, but qRT-PCR confirmed that FR235222-treatment significantly upregulated expression of *TA05055, TA13515* and *TA16485* (**Supplementary Fig. 7b**). Transcription factor binding sites have been defined for *TA11145* (Taap2.me1) and PF3D7_1466400 (PfAP2-EXP) the orthologue of *TA08375*^21, 38^. So, using the motif search function at PiroplasmaDB we asked if their respective binding sites are located within 1000bp upstream of each of the 441 DEG genes. This revealed that upregulation of TaAP2-EXP could be potentially responsible for driving transcription of 93 DEGs and upregulation of Taap2.me1 for 68 DEGs (**Supplementary File 1**). Only 13 DEGs harbored binding sites for both AP2s and when taken together FR235222-induced upregulation of Taap2.me1 (*TA11145*) and TaAP2-EXP (*TA08375*) could be responsible for increased expression of 33% of the 441 DEG. However, 3 additional ApiAP2 genes are also upregulated by FR235222 and they might be responsible for driving expression of the remaining 67% of DEGs. Clearly, blocking TaHDAC1 activity has the potential to initiate a cascade of transcription factor activity that promotes commitment to merogony. Moreover, it implies that when active TaHDAC1 represses schizont expression of these merogony-associated ApiAP2s and taken together, this places TaHDAC1 as major transcriptional modulator of merogony by controlling expression of a subset *T. annulata* ApiAP2 transcription factors and their target genes.

**Table 3.**
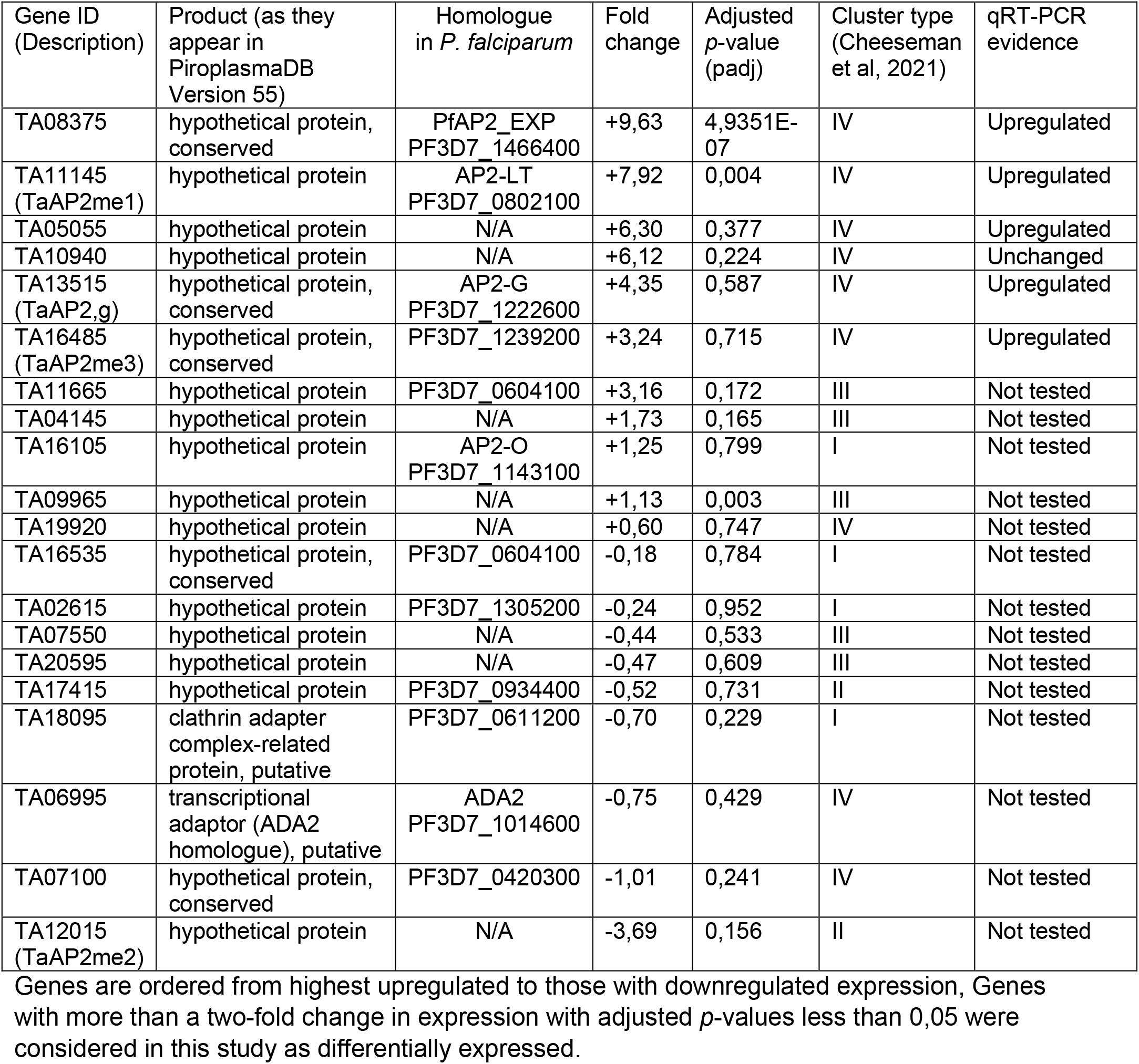
Summary of RNA-seq and qRT-PCR results for predicted *Theileria annulata* proteins with ApiAP2 transcription factor domains (PF00847).

## Discussion

In this study, we observed that FR235222 treatment of *T. annulata* schizonts led to significant deregulation of 468 genes and one likely reason for this small number compared to the total number of 3796 predicted genes is that FR235222 is specific for a single parasite HDAC (TaHDAC1). The *T. annulata* RNA-seq data was generated following 48 h of FR235222 treatment, a timepoint when treatment had no effect on proliferation of non-infected BL20 cells. In spite of no known pronounced merozoite-associated dampening of host cell proliferation at 48 h some parasite genes classically associated with merozoite invasion of red blood cells became actively transcribed e.g. (TA05705, TA05815, TA16660 and TA02980=TaAMA1) coding for rhoptry and microneme proteins. This argues that FR235222-mediated inhibition of TaHDAC1 had triggered a switch in gene expression enabling schizonts to progress toward merogony. *In vitro* merogony in *T. annulata* takes 7-8 days to complete and a temporal coordination of schizont to merozoite gene expression has been proposed^39^. Comparison of our 48 h FR235222-generated RNA-seq data to that obtained from proliferation arrested merozoites^21^ identified only 24 common genes (**Supplementary File 1**) among which are homologs of PfTRAP, PfRON5, PfRON11 and a glideosome-associated connector, which suggests that their induction is linked to initiation of merogony. Seven days of treatment with FR235222 resulted in a complete arrest in host/parasite proliferation and expression of merozoite-specific rhoptry protein (Tamr1) with rhoptry formation an indication of late stage merogony^40, 41^. In addition, inhibition of TaHDAC1 for 7 days led to an increase in the number of cells displaying small condensed parasite nuclei representative of merozoite production: reinforcing the conclusion that FR235222-treatment promotes hyperacetylation of chromatin associated with a subset of genes, whose elevated transcription is required to initiate the schizont to merozoite transition.

Recently, four anti-cancer HDAC inhibitors were tested against *T. annulata*-transformed leukocytes and an irreversible arrest in proliferation was observed and ascribed to parasite death and host cell apoptosis^42^. We confirmed that apicidin enhances merogony induced by elevated temperature (41°C), but not at 37°C, and it did not inhibit down regulation of the macroschizont marker (p104) in cells undergoing merogony suggesting initial down regulation of this schizont marker does not occur via histone deacetylation, as also indicated for FR235222 (see Fig 2d). By contrast, at the 10 nM dose used the Apicomplexa HDAC1-specific inhibitor FR235222 had no effect on host leukocyte viability and arrested host cell proliferation was due to formation of merozoites. These differences are most easily explained by apicidin and other anti-cancer drugs inhibiting both host cell and parasite HDAC activities, but clearly 7-days culture at 41°C is a stronger inducer of merogony than 7-days of unique FR235222 treatment at 37°C.

ApiAP2 transcription factors are known to operate in transcriptional cascades to affect changes in expression of sets of parasite genes^43, 44^. In addition to three previously described merozoite-associated ApiAP2s, FR235222 mediated upregulation involves several other *T. annulata* ApiAP2s including TA08375. The AP2 domain of TA08375 displays significant similarity to PfAP2-EXP that in *P. falciparum* regulates expression of multigene families like *stevors* and *rifins*, whose corresponding proteins are secreted into the red blood cell^38^. Another significantly upregulated ApiAP2 following 7 days of FR235222 treatment was TA11145 (*Taap2*.*me1*) previously identified as an ApiAP2 involved in merogony^21^. Thus, in *T. annulata* schizonts the level of acetylation mediated by TaHDAC1 appears to function as an epigenetic repressor of these ApiAP2 genes that become de-repressed in merozoites. It’s noteworthy that all FR235222-induced ApiAP2s are cluster IV genes that are repressed in schizonts by monomethylation of H3K18^27^.

In summary, lack of tools to genetically modify *Theileria* parasites led us to take a chemical genetic approach to knockout TaHDAC1 activity and transcriptionally and phenotypically profile FR235222-treated schizonts. Our results when combined with those of Cheeseman et al.^27^ argue that in *Theileria annulata* expression/repression of merozoite genes involves a balance of histone acetylation versus methylation and that modulation of this balance by pharmacological inhibition of deacetylases and/or methyltransferases impacts on the parasite’s developmental fate.

## Methods

### Cell culture

Virulent *Theileria annulata*-transformed Ode macrophages (first isolated in Anand India) at passage 52^20^ were cultured in RPMI that contained 10% FBS (Gibco), penicillin/streptomycin (Gibco), HEPES (Euromedex) and L-glutamine. The D7 cell line^45^, bovine leukemia virus immortalized B cells (BL20)^46^ and their *T. annulata*-transformed counterparts (TBL20, Hissar parasite strain)^47^ were cultured under the same conditions except that 2-mercaptoethanol (Gibco) was added to the culture medium. Cultures were passaged 3-times per week.

### Immunostaining of cells

Cells in suspension were first washed with PBS (LONZA) and cell numbers adjusted to a concentration of 10^5^ ml^−1^ in PBS. Cells were then adhered to SuperFrost™ microscopic slides (Thermo Scientific) by performing cytospin (Cellspin® II, 3 minutes at 1500 rpm). Schizonts purified from TBL20 cells were diluted in PBS and attached to poly-L-lysine coated glass cover slips (Corning® BioCoat®). Fixation was done with 4% paraformaldehyde (PFA, Electron Microscopy Sciences, Hatfield, PA) solution for 10 mins at room temperature. Primary antibodies were: rabbit polyclonal antibody against Tamr1 (mAb 1D11), anti-Ta-p104 (Clone IC12, polyclonal mouse antiserum) and rabbit monoclonal anti-acetyl H4 (Sigma-Aldrich). Antibodies were diluted in PBS-1% BSA-0.3% Triton X100 (PBST). Secondary antibodies were a goat anti-mouse Alexa Fluor® 488 and a goat anti-rabbit Alexa Fluor® 594 (Invitrogen™) diluted at 1:2000 in PBS containing 0.3% Triton X100. Cells were finally covered by a round coverslip and sealed using Invitrogen™ ProLong™ Gold antifade mountant with DAPI (ThermoFisher).

### Hemacolor® staining of leukocytes

Following cytospin of leukocytes the Hemacolor® staining kit (Merck, Germany) that works based on eosin (red)/azur (blue) staining was used. Microscopic slides were observed under a normal light microscope (Leica DM750).

### Parasite purification from transformed leukocytes

*T. annulata* schizonts were purified using a previously described method except that aprotinin was omitted from the protocol^48^. Finally, the schizont fraction was purified from whole cell components through Nycodenz (Axon) density gradient centrifugation. 40-50×10^6^ infected leukocytes were used for each purification.

### Western blotting

Cells were lysed using X1 RIPA buffer (ChromoTek) containing protease and phosphatase inhibitors (100X, Halt™ protease and phosphatase inhibitor cocktail, Thermo Scientific). Protein concentration of lysates was measured by Bradford assay. Lysates were mixed with 4X Laemmli buffer (BIO-RAD) and boiled for 5 mins at 95°C. Next, equal amounts of lysate were loaded into wells of 12% Mini-PROTEAN® TGX 4-20% pre-cast protein gel (BIO-RAD) installed in Mini-PROTEAN® tetra vertical electrophoresis cell filled with 1X TG-SDS solution (Euromedex). The gels were run for 1 h at constant 150 volts. Next, proteins were transferred to nitrocellulose membranes using iBlot™ gel transfer device (ThermoFisher) program P0. Following protein transfer, filters were blocked for 1 h at room temperature in PBS solution containing 5% W/V dry milk (Régilait, France). Filters were incubated in a cold room (4°C) overnight with primary antibodies diluted in PBS with 0.1% TWEEN® 20 (Merck) solution (1:1000 dilution). Secondary antibodies (GeneTex) were diluted in the same solution (1:5000) and were incubated with the membrane for 1 h at room temperature. Finally, protein bands were rendered visible by adding Pierce™ ECL western blotting substrate (Thermo Scientific) on the filters and images were taken by Fusion FX western blot imaging machine (Vilber Lourmat). Mouse monoclonal anti-α-tubulin (Sigma-Aldrich®) antibody was used as loading control.

### Quantitative real time PCR (qRT-PCR) and primers

Cells were washed in PBS and total RNA was extracted using RNAeasy® plus mini kit (QIAGEN). The quality and quantity of extracted RNA was evaluated by a NanoDrop 1000 machine (ThermoFisher). Complementary DNA (cDNA) synthesis was done by using 1 μg RNA as template and the Moloney Murine Leukemia Virus (M-MLV) reverse transcriptase (Promega), in 20 μl reaction volume according to the instructions manual. To perform qRT-PCR target sequences were amplified in a SYBR green PCR master mix (ThermoFisher Scientific) mixed with diluted cDNA (1:20), double distilled water and primers. The PCR was run in LightCycler® 480 instrument (Roche) and results analysed by dedicated software. Finally, the 2^−ΔΔCT^ methodology was employed to estimate relative gene expression levels^49^. *T. annulata* heat shock protein 70 (TaHSP70=TA11610) was used as the house keeping gene for normalization. The list of qPCR primers is provided in **Supplementary File 2**.

### Confocal microscopy and preparation of images

Immunolabelled cells were studied either under a confocal SP8 laser microscope (Zeiss), or a Leica DMi8 epifluorescence microscope. Images were taken by LASX software and analysed by ImageJ.

### Manual counting of schizont nuclei and Tamr1+ cells

For better visualization and quantification of parasite nuclei per infected leukocyte under different experimental conditions, cells were first well spread on microscopic slides by cytospin. The cytospinned cells were then fixed by 4% PFA solution and their DNA stained with DAPI dye (ProLong™ Gold antifade mountant with DAPI, ThermoFisher). The number of parasite nuclei per infected leukocyte were then counted manually in 50 infected leukocytes under an epifluorescence microscope. Similarly, Tamr1 expressing infected leukocytes were quantified in cytospinned immunostained slides.

### Measurement of leukocyte proliferation and viability

An automatic cell-counting machine (BIO-RAD TC20) that functions based on the trypan blue exclusion method was used to measure cell numbers. Samples were prepared according to manufacturer’s instructions.

### RNA preparation, sequencing and bioinformatics analyses

RNAeasy Plus Mini Kit (Qiagen) was used to extract total RNA from TBL20 cells treated or not with 10nM FR235222 for 48 h. Samples were prepared in triplicate and contained 4.5-20 μg RNA per sample. Desiccated samples (RNAstable®, Biomatrica®) were shipped to the Pathogen Genomics Laboratory, King Abdullah University of Science and Technology (KAUST) in Saudi Arabia. Upon arrival an Agilent RNA 6000 Nano kit and Qubit Broad Range kit were utilized to check the quality and quantity of the RNA. The RNA libraries were prepared according to the Illumina TruSeq RNA Sample Preparation protocol. Libraries were then processed for deep, paired-end (2 × 150 bp) sequencing with the Hiseq 4000 platform (Illumina, USA). The resulting FASTQ files were then aligned against the reference genome (Ankara C9 genome). Reads were counted and genes with less that 5 reads per triplicate samples were eliminated. Next, the counting data was analysed by a DESeq2 pipeline^50^ to determine the proportion of differentially expressed genes between control and treated samples. Data normalization was made according to the size factor per sample (i.e. geometric mean per gene and the median that gives a size factor that does not include genes at zero). Wald test was used to compare the two groups and to calculate p values taking into account the heterogeneity for the padj (if too heterogeneous then NA). In the current study we considered genes displaying more than 2 Log2 fold change in expression with an adjusted *p*-value (padj) less than 0.05 as differentially expressed (DE). RNA-Seq reads have been uploaded to the European Nucleotide Archive (https://www.ebi.ac.uk/ena/) under the Study accession number PRJEB50801.

## Data availability

RNA-Seq reads that support the findings of this study have been deposited in European Nucleotide Archive (ENA) with the accession code PRJEB50801. Authors confirm that all other relevant data are included in the paper and/ or its supplementary information files.

## Acknowledgments

ST acknowledges a LabEx ParaFrap postdoctoral fellowship and GL acknowledges ANR-11-LABX-0024 support and core funding from INSERM and the CNRS. AC received funding from Head of College Scholars List Scheme, College of Medical, Veterinary and Life Sciences, University of Glasgow for a summer project. AP acknowledges the financial support received from KAUST in form of OCRF-20146CRG4 and BAS/1/1020-01-01 grants. We thank members of the KAUST Bioscience Core Laboratory (BCL) to generate the raw sequence datasets. We thank

## Author Contributions

S.T., L.M., S.M. and A.C. performed experiments.

S.T., B.S., A.C., T.M., B.R.S., F.A., A.P. and G.L. analysed data.

S.T. and G.L. designed the study.

B.R.S., F.A., A.P., and G.L. contributed essential materials.

S.T. and G.L. wrote the manuscript.

All authors read and approved the manuscript.

## Competing interests statement

The authors declare no competing interests.

## Additional Information

Supplementary Information is available for this paper.

Correspondence and requests for materials should be addressed to Gordon Langsley. Reprints and permissions information is available at www.nature.com/reprints

## Supplementary Figures

**Fig. S1.**
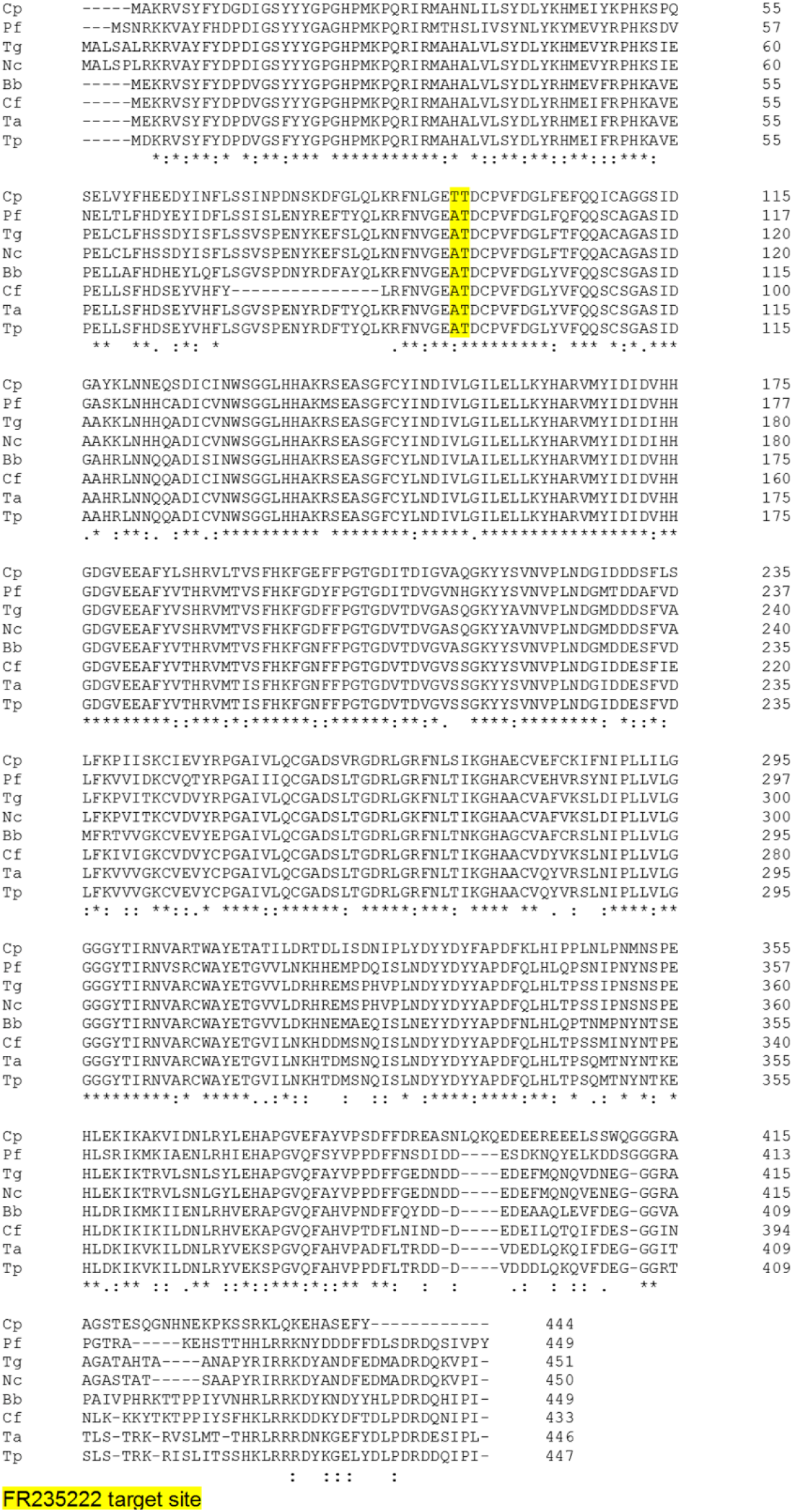
Genomes of medical and veterinary important apicomplexan genera possess a highly conserved histone deacetylase (HDAC1). Protein sequence alignment of HDAC1 from several apicomplexan genera. Bb, *Babesia bovis*; Cf, *Cytauxzoon felis*; Cp, *Cryptosporidium parvum*; Nc, *Neospora caninum*; Pf, *Plasmodium falciparum*; Ta, *Theileria annulata*; Tp, *Theileria parva*; Tg, *Toxoplasma gondii*. Multiple sequence alignment was done by Clustal Omega online (https://www.ebi.ac.uk/Tools/msa/clustalo/).

**Fig. S2.**
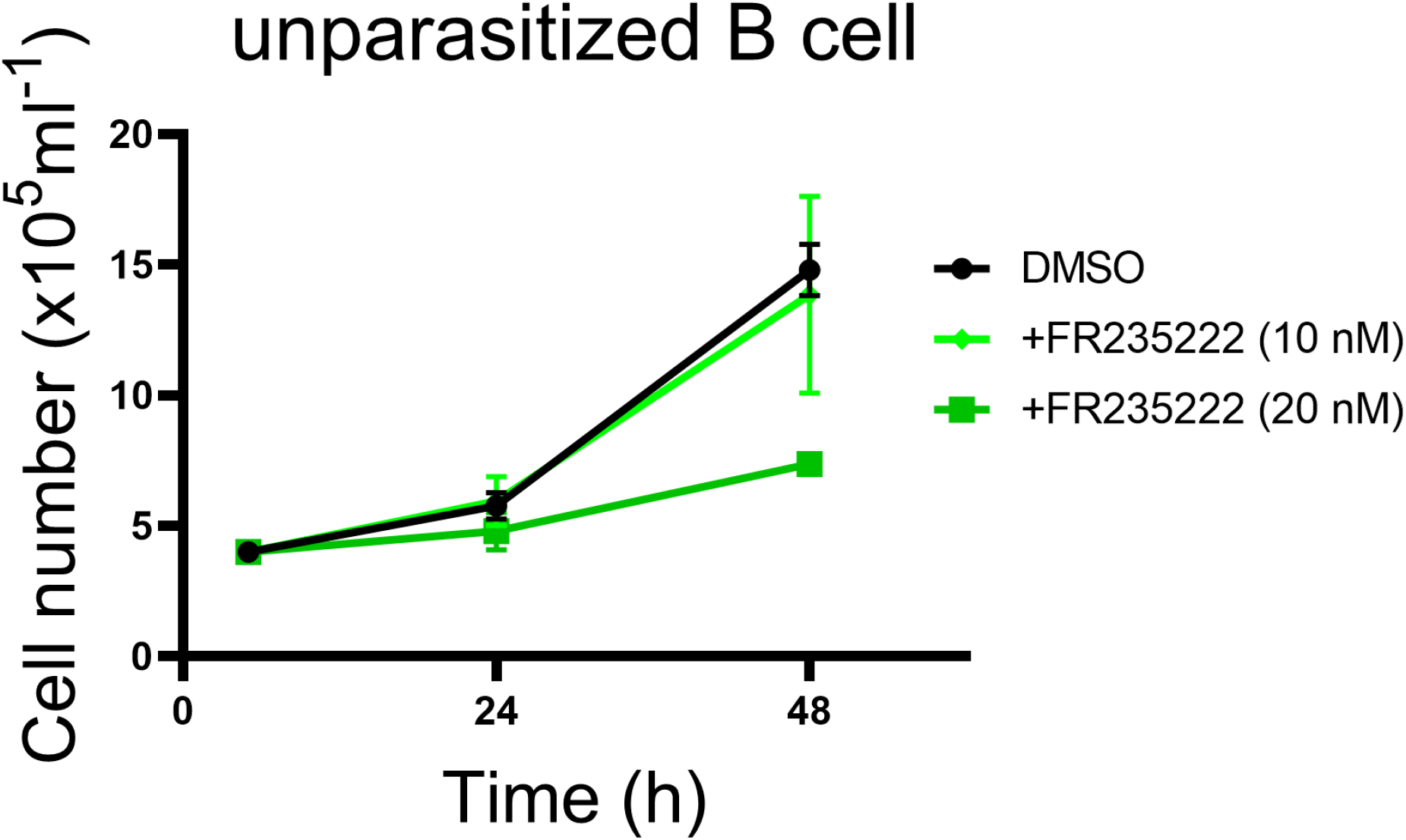
Proliferation of non-infected bovine B cells (BL20) undergoing FR235222 treatment. BL20 B cells constantly proliferate, as they are immortalized (but not transformed) by bovine leukemia virus (BLV).

**Fig. S3.**
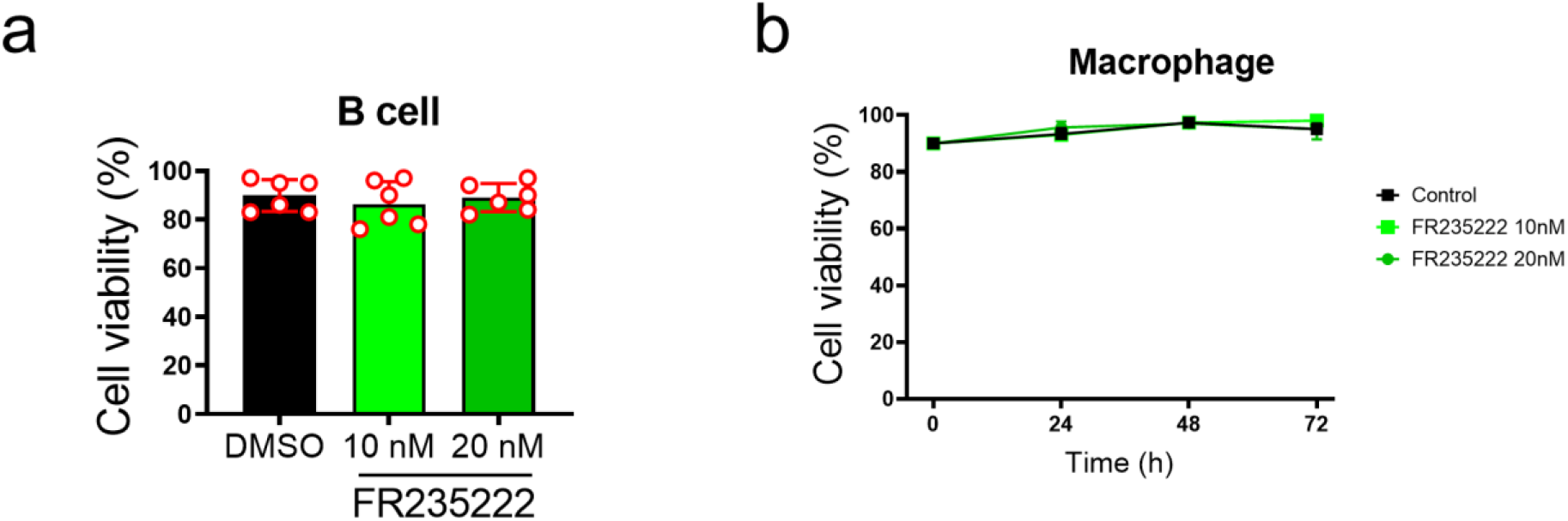
FR235222 treatment and cell viability of *Theileria*-transformed leukocytes. **a** Viability measurement of TBL20 B cells post 48 h exposure to 10 and 20 nM FR235222 compared to DMSO-only control TBL20. **b** Viability of virulent Ode macrophages (p52) throughout 72 h treatment with FR235222 TaHDAC1 inhibitor. Each graph is representative of several independent experiments.

**Fig. S4.**
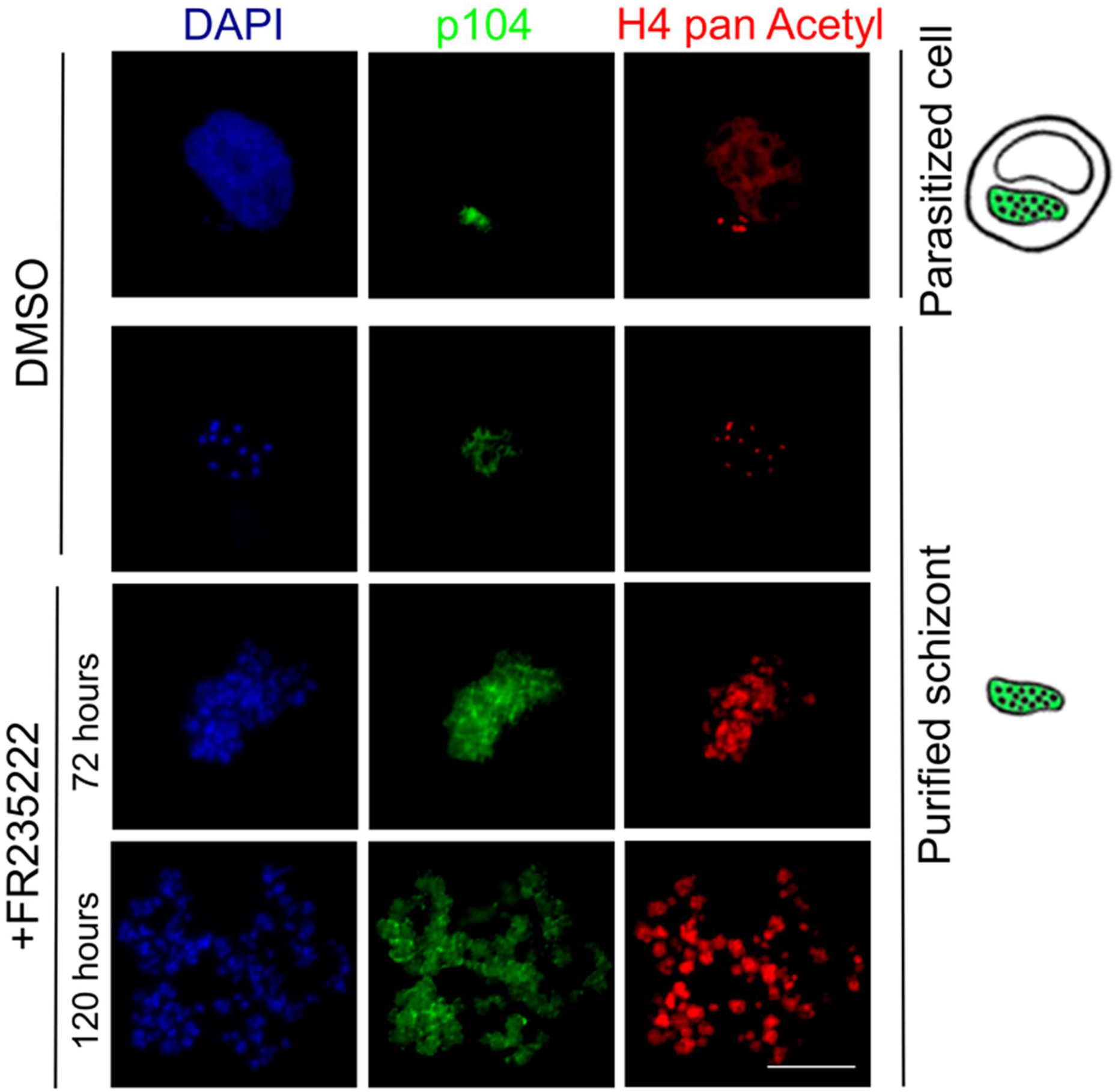
Hyperacetylation of *Theileria annulata* histone 4 following TaHDAC1 inhibition by FR235222. Immunofluorescence images of TBL20 B cells treated with FR235222. Top row shows a non-treated control. The other rows show parasites purified from treated and control cultures. Parasites were decorated with the 1C12 monoclonal antibody to p104. A rabbit monoclonal antibody detects acetylation of histone 4 at K5, K8 and K12 residues. Host and parasite nuclei stained with DAPI dye. Note the increase in parasite nuclei and H4 acetylation upon FR235222 exposure. X100 magnification, scale bar = 10 μm.

**Fig. S5.**
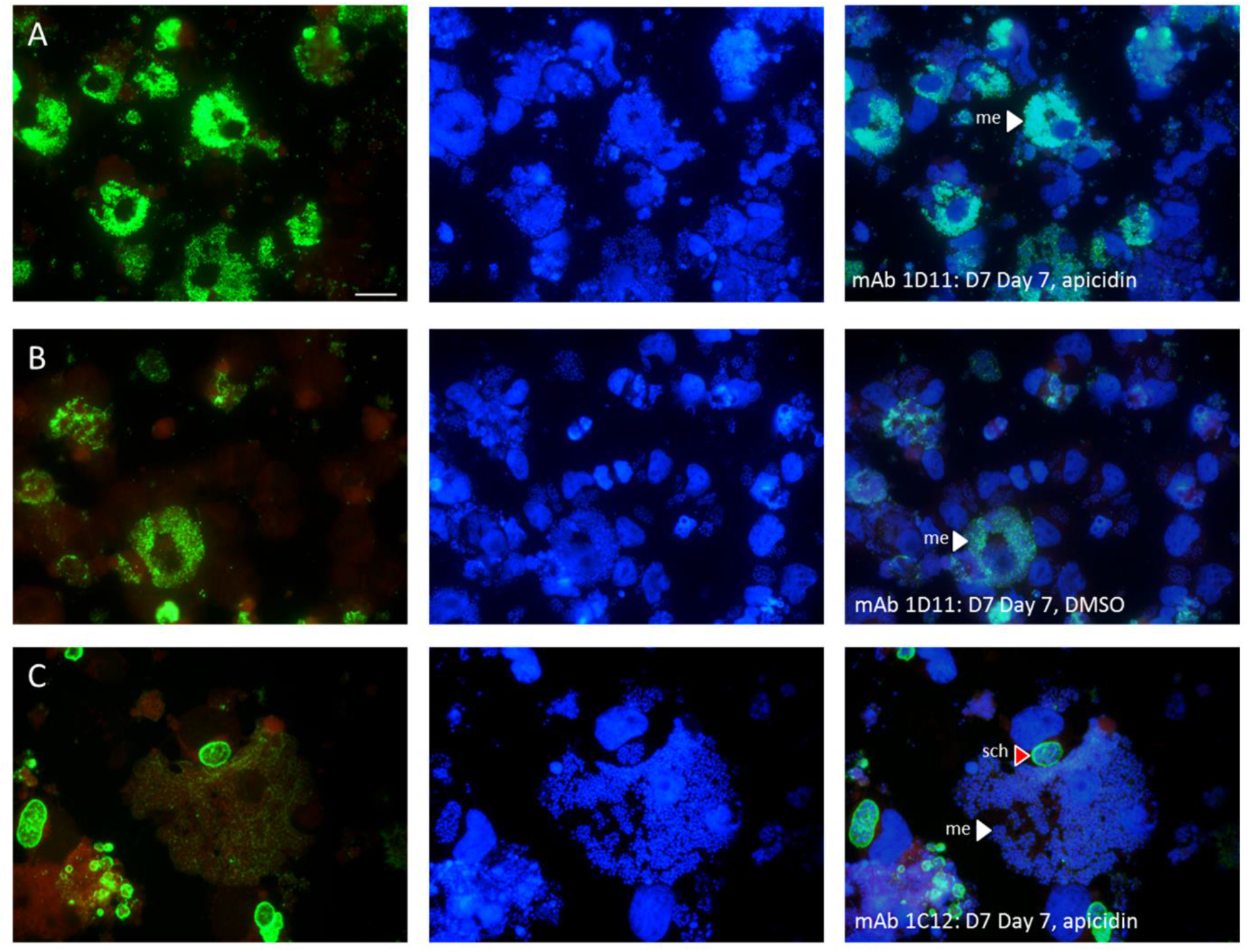
Indirect immunofluorescence using mAb 1D11 that detects merozoite antigen Tamr1 or mAb 1C12 that detects the macroschizont p104 antigen. a and c row represent cells derived from culture incubated with apicidin 25 nM; b row DMSO control culture. Left column is green image of mAb antibody staining; middle shows DAPI stained nuclei; right column is a merge of first two images. White arrow heads denote cells undergoing merogony, red arrow head denotes macroschizont, scale bar = 10 μm.

**Fig. S6.**
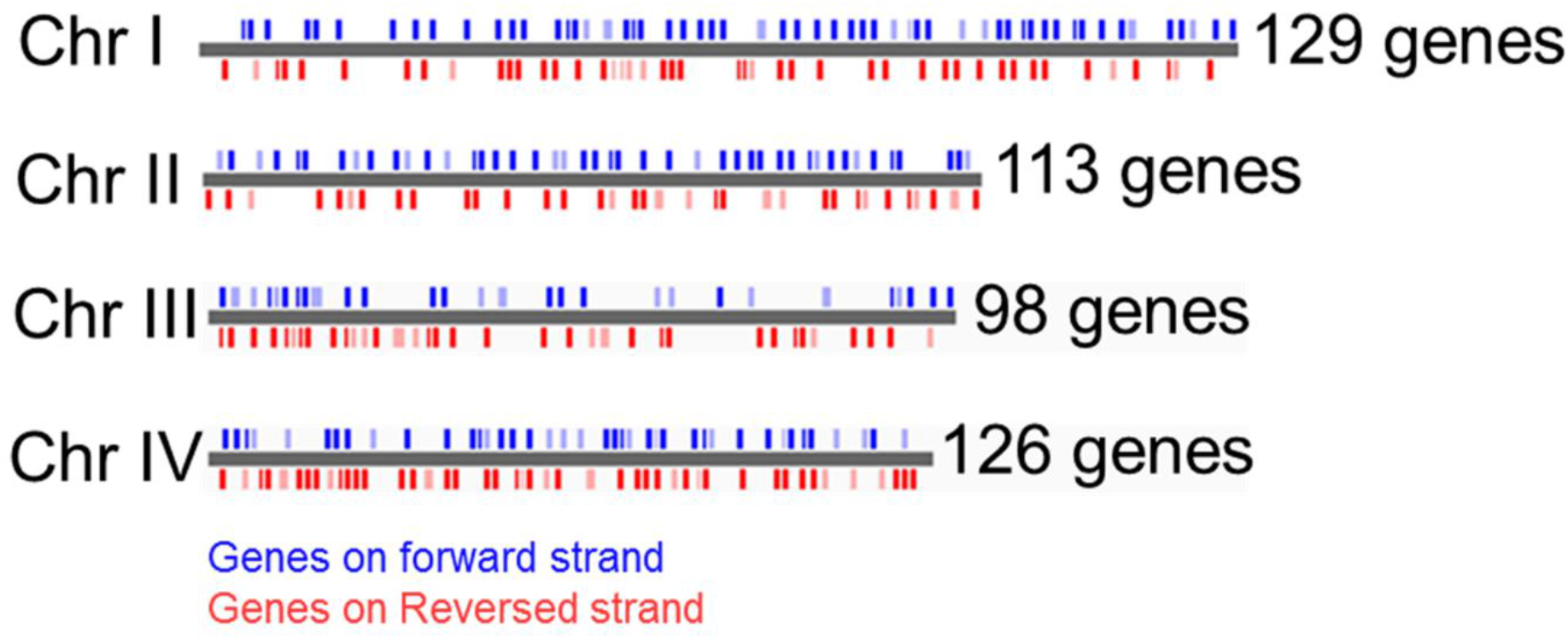
Genomic positioning of 468 (441 up- plus 27 downregulated DEG genes) FR235222-induced differentially expressed genes positioned across the four *T. annulata* chromosomes. Image generated at PiroplasmaDB (https://piroplasmadb.org/piro/app) using the genome viewer feature. Note that two DEGs are mitochondrial and thus not shown.

**Fig. S7.**
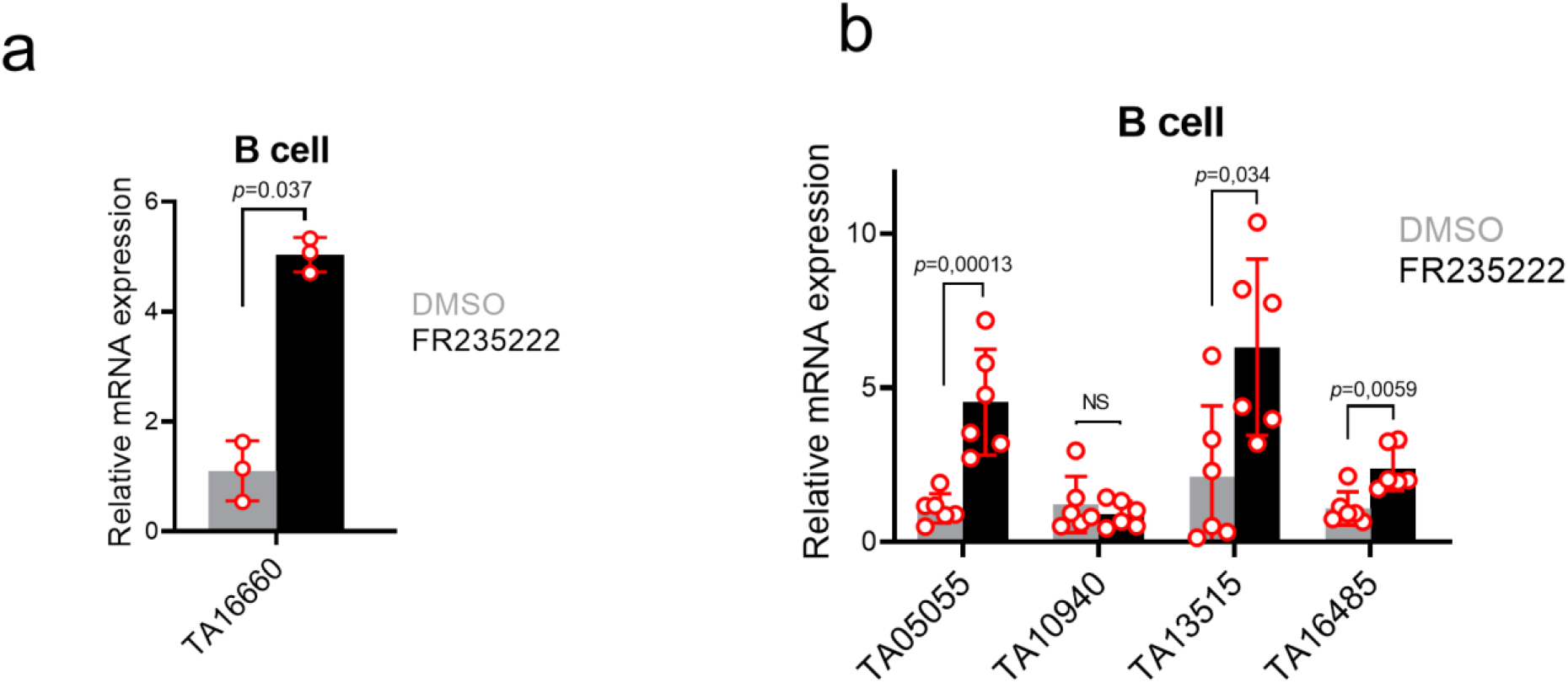
FR235222-induction of *T. annulata* genes. qRT-PCR results on expression of TA16660 (TaRON) **a** and some AP2 genes **b** in TBL20 cells. Based on RNA-seq data the AP2 genes displayed >3-fold augmentation in expression with non-significant adjusted *p*-values (See Table 3). Results are representative of two biological replicates. *p* values were determined by Student’s two-tailed t test. NS: not significant. Error bars depict standard deviations.

**Supplementary Table 1:** List of *Theileria annulata* genes verified by qRT-PCR and presented in Figure 2a. FR235222-target genes in *Toxoplasma gondii* were identified by Bougdour et al, (2009).

| Gene category | Gene ID | Expressed sequence tag | *Toxoplasma* orthologue ID |
| --- | --- | --- | --- |
| ***Theileria annulata***<br>**orthologues of**<br>***Toxoplasma gondii***<br>**FR235222 target genes**<br>(n=13) | TA03055 | Macroschizont | TGME49_230460 |
|  | TA05900 | No EST | TGME49_309980 |
|  | TA06500 | Merozoite, Piroplasm | TGME49_308960 |
|  | TA07295 | Piroplasm | TGME49_294600 |
|  | TA07655 | Macroschizont | TGME49_211430 |
|  | TA11375 | Macroschizont, Merozoite | TGME49_319988 |
|  | TA11395 | No EST | TGME49_275360 |
|  | TA15060 | Macroschizont, Merozoite, Piroplasm | TGME49_226310 |
|  | TA16180 | No EST | TGME49_205750 |
|  | TA16310 | Merozoite | TGME49_230020 |
|  | TA16570 | Merozoite, Piroplasm | TGME49_242110 |
|  | TA17985 | No EST | TGME49_206650 |
|  | TA19895 | Macroschizont | TGME49_265542 |
| **Random genes**<br>(n=17) | TA02685 | Macroschizont, Piroplasm |  |
|  | TA02980 | Merozoite |  |
|  | TA04560 | Macroschizont |  |
|  | TA06655 | Macroschizont, Merozoite |  |
|  | TA07550 | Merozoite, Piroplasm |  |
|  | TA08425 | Macroschizont, Merozoite |  |
|  | TA09495 | No EST |  |
|  | TA09590 | Macroschizont |  |
|  | TA09760 | Macroschizont |  |
|  | TA09995 | No EST |  |
|  | TA10915 | Macroschizont, Merozoite |  |
|  | TA11405 | Macroschizont |  |
|  | TA13175 | Macroschizont |  |
|  | TA16685 | Merozoite |  |
|  | TA18055 | Macroschizont, Merozoite, Piroplasm |  |
|  | TA19600 | Macroschizont, Merozoite |  |
|  | TA19860 | Macroschizont |  |

**Supplementary Table 2:** Effect of apicidin treatment on parasite differentiation potential of infected cloned cell line D7 at 37°C and 41°C.

| **Anti-Tamr1 +ve cells in drug treated Day 7 37°C cultures** |  |  |
| --- | --- | --- |
| Treatment | % +ve | Fold increase (Apicidin/DMSO) |
| Apicidin 25nM | 0 | 0 |
| DMSO | 0 |  |
| **Anti-Tamr1 +ve cells in apicidin treated Vs DMSO 41°C Day 7 cultures** |  |  |
| Treatment | % +ve | Fold increase (Apicidin/DMSO) |
| Apicidin 25nM | 28.4* | 1.44 |
| DMSO | 19.7 |  |

\* Significant difference between drug and control culture (P<0.01).
Immunofluorescence was performed as described in<sup>40</sup>; quantification of differentiation and statistical analysis using $\chi^2$ test was as described in<sup>41</sup>.

## Notes

https://www.ebi.ac.uk/ena/browser/view/PRJEB50801

### Competing Interest Statement

The authors have declared no competing interest.

